# *Max* deletion destabilizes MYC protein and abrogates Eμ-*Myc* lymphomagenesis

**DOI:** 10.1101/556522

**Authors:** Haritha Mathsyaraja, Brian Freie, Pei-Feng Cheng, Ekaterina Babaeva, Derek Janssens, Steven Henikoff, Robert N Eisenman

**Affiliations:** Basic Sciences Division, Fred Hutchinson Cancer Research Center, Seattle, WA

## Abstract

Although MAX is widely regarded as an obligate dimerization partner for MYC, its function in normal development and neoplasia is not well defined. We show that B-cell specific deletion of *Max* has a surprisingly modest effect on B-cell development but completely abrogates Eµ-*Myc* driven lymphomagenesis. In both contexts, MAX loss leads to a significant reduction in MYC protein levels. This outcome is associated with the downregulation of numerous transcriptional targets of MAX including a subset that regulate MYC stability. Reduction in MYC protein levels is also observed in multiple cell lines treated with a MYC-MAX dimerization inhibitor. Our work uncovers a layer of *Myc* autoregulation critical for lymphomagenesis yet partly dispensable for normal lymphoid development.

## INTRODUCTION

The MAX protein was first identified as a specific dimerization partner with members of the MYC oncoprotein family (MYC, MYCN and MYCL). Like MYC, MAX is a member of the basic region-helix-loop-helix-zipper (bHLHZ) class of transcriptional regulators, and the association of MYC with MAX is mediated by heterodimerization between their two HLHZ domains. MYC-MAX heterodimers bind DNA through direct contact of each bHLHZ basic region with the major groove of E-box DNA sequences (CANNTG) (for reviews see (Conacci-Sorrell et al. 2014) (Carroll et al. 2018)). MYC does not homodimerize or bind DNA under physiological conditions and, aside from MAX, no other bHLHZ proteins have been compellingly demonstrated to dimerize with MYC. Because mutations in the MYC bHLHZ that prevent association with MAX also block MYC’s major biological activities, it has been generally assumed that MAX is required for MYC function. Moreover, targeted deletion of *Max* in mice results in early post-implantation lethality, consistent with essential functions for *Myc* and *MycN* during embryonic development (Shen-li et al. 2000). In addition to dimerizing with MYC family proteins, MAX also forms E-box DNA binding heterodimers with the MXD family, MNT and MGA proteins, all of which act as transcriptional repressors.

Despite the apparent centrality of MAX for the functions of multiple bHLHZ transcription factors there is evidence that MAX loss of function can be tolerated and even oncogenic in several biological contexts. For example, pheochromocytoma cell lines can proliferate in the absence of MAX and a subset of familial pheochromocytomas are strongly associated with inactivation of *Max* (Hopewell and Ziff 1995) (Comino-Méndez et al. 2011). In addition, approximately 6% of human small cell lung carcinomas (SCLC) exhibit loss of MAX, and introduction of MAX into human SCLC lines lacking MAX arrests growth (Romero et al. 2014). Lastly, in *Drosophila melanogaster* larval development is less compromised by loss of MAX than by loss of MYC, and several critical activities of MYC appear unaffected by MAX inactivation (Steiger et al. 2008). These findings suggest that there are functions of MYC independent of MAX and that loss of MAX in some settings can promote oncogenic conversion.

To investigate a MAX-independent role in MYC induced oncogenesis we turned to Eμ-*Myc* transgenic mice which model the 8;14 translocation found in Burkitt’s B cell lymphomas and have provided many insights into MYC driven lymphomagenesis. The over-expression of MYC produces a polyclonal increase in pre-B cells in young mice, accompanied by reduced differentiation to mature B cells (Harris et al. 1988). Earlier work, using an Eµ-*Max* transgene, established that overexpression of MAX alone in murine lymphoid cells is non-oncogenic and results in reduced B-cell proliferation and numbers. Importantly, in the context of an Eµ-*Myc* transgene, augmented expression of *Max* also attenuated B cell lymphomagenesis and reduced lymphoproliferation (Lindeman et al. 1995), indicating that the ratio of MYC:MAX expression levels can influence MYC function. However, the requirement for endogenous MAX in MYC induced tumorigenesis has not been determined. To address these questions, we generated a conditional *Max* allele to elucidate *Max* function in lymphomagenesis and in B-cell homeostasis.

## RESULTS AND DISCUSSION

### ***Max* deletion partially impairs B cell development**

We constructed a *Max* targeting vector by inserting *loxP* sites (and a Neomycin resistance gene) flanking exon 4 within a full-length *Max* genomic clone. This region encodes nearly the entire helix 2-leucine zipper region of *Max* necessary for dimerization with MYC and other bHLHZ proteins (Fig. 1A), and its Cre-mediated deletion is expected to lead to a frame shift and truncation within exon 5, leading to a 127aa protein lacking the HLHZ domain. Expression of Cre in *Max*^fl/+^ ES cells resulted in heterozygous deletion of *Max* (*Max*^Δ/+^) and these ES cells were employed to produce chimeric mice. Extensive intercrossing of *Max*^Δ/+^ F1 mice failed to produce any homozygous *Max* null offspring, consistent with a previous report (Supplemental Fig. S1A) (Shen-li et al. 2000).

**Fig. 1:**
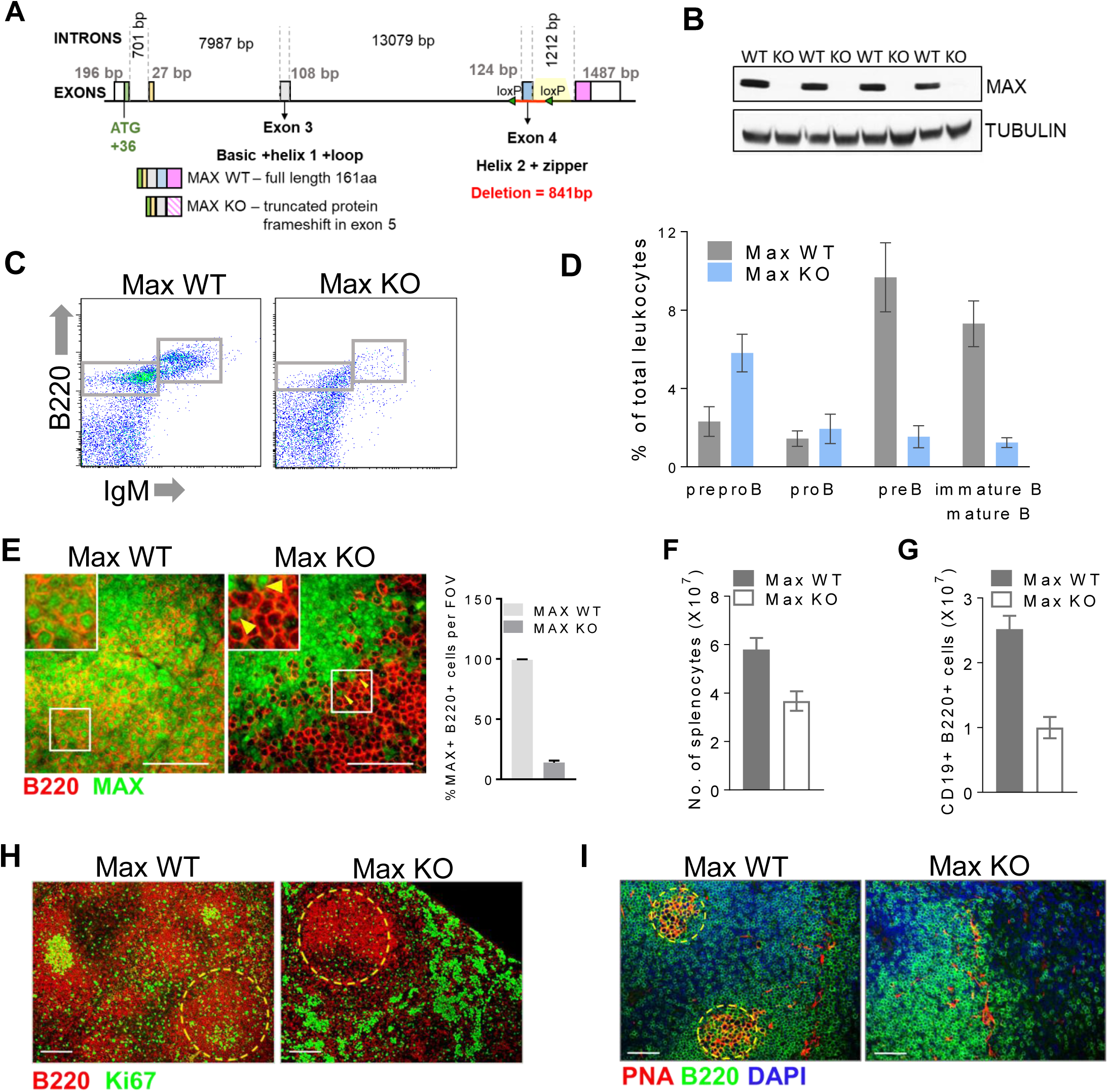
Conditional Deletion of *Max* in the B-cell Lineage. (A) Schematic depicting location of loxP sites at the *Max* locus. (B) Representative immunoblots for MAX in B220+ splenocytes from *Max* fl/fl (WT) and *Max* fl/fl MBI-cre (KO) animals. (C) Representative flow plots showing B220+ and IgM+ populations in CD45 gated BM cells. (D) Quantification of B lymphocyte precursor populations in *Max* WT (n=5) and KO (n=6) bone marrow. (E) Dual immunofluorescence (IF) for MAX and B220 in spleens. Quantification of MAX+ B220+ cells from MAX KO spleens (n=3). Yellow arrows indicate MAX+B220+ cells in Max KO. Total number of (F) splenocytes and (G) CD19+ B220+ cells in *Max* WT and KO mice (WT n=8 KO n=9). (H) IF staining for B220 and proliferation marker Ki67 in *Max* WT and KO spleens. (I) Micrographs showing germinal centers (PNA) in *Max* WT and KO spleens (representative image n=3 animals per genotype). Scale bars = 100µM.

We next crossed *Max*^fl/fl^ mice with hemizygous mb1-Cre transgenic mice. Expression of the mb1 gene is restricted to B-cells beginning at the early pro-B-cell stage and mb1-Cre has been used extensively to study B-cell development and function (Hobeika et al. 2006). Immunoprecipitation from B-cell lysates derived from *Max*^fl/fl^, and mb1-Cre; *Max*^fl/fl^ mice using an anti-C-terminal MAX serum detected the MAX protein band in *Max*^fl/fl^ mice (referred to herein as *Max* WT) while mb1-Cre; *Max*^fl/fl^ cells (referred to as *Max* KO) did not express any protein reactive with the antibody (Fig. 1B) as expected upon Cre-mediated deletion.

We examined the consequences of *Max* deletion on normal B-cell development by comparing *Max* WT with *Max* KO mice. Using flow cytometry to assess cell sub-populations in the B-cell lineage, we noted a significant decrease in the numbers of B220 positive, IgM^-^ and IgM^+^ B-cells from Max KO relative to wildtype (Fig. 1C). Notably B220+ IgM+ B-cells (pre-B-cells) were nearly 10-folder lower in *Max* KO samples than in *Max* WT (Fig. 1D). More detailed analysis of different stages of B-cell development showed that while the proportion of prepro- and pro-B-cells were approximately the same in mice of the two genotypes, the percentage of pre-B, immature-B and mature B-cells was strikingly diminished in *Max* KO mice, indicating loss of *Max* results in a significant block in pro-B to pre-B-cell differentiation (Fig. 1D). This block in development is similar to that seen upon *Myc* loss in B cells (Habib et al, 2007). In addition, BM precursors from *Max* KO mice failed to efficiently differentiate into B220+ cells upon treatment with IL-7 *in vitro* (Supplemental Fig S1B, C). We also noted a compensatory increase in the percentages of CD3+ T cells and CD11b+ myeloid cells (Supplemental Fig. S1D). To study mature B-cell populations, we examined spleens of *Max* KO mice. A majority (∼86%) of B220+ cells lacked detectable MAX staining in their nuclei (Fig. 1E). This correlated with reduced numbers of total splenocytes (Fig.1F) and CD19+ B220+ splenocytes in the *Max* KO spleens compared to *Max* WT (Fig.1G, Supplemental Fig. S1E). Indeed, B220+ areas in the spleen displayed a significant reduction in Ki67 staining (Fig. 1H, Supplemental Fig. S1F).

Intriguingly, loss of Ki67 appeared to be most dramatic in areas corresponding to germinal centers (GC). Since MYC is known to play a critical role in GC formation and maintenance (Calado et al. 2012), we stained spleens for PNA, a GC B-cell marker. Although non-immunized mice have relatively few GCs, we still observed positive staining for PNA in *Max* WT mice whereas *Max* KO spleens exhibited strongly reduced PNA staining (Fig 1I), suggesting MAX plays a critical role in GC formation. Our results suggest that *Max* is not essential for B lymphocyte development and differentiation, a conclusion in agreement with the recent study by Perez-Olivares et al. examining the loss of *Max* in mouse B-cells (Perez-Olivares et al. 2018).

Similarly, depletion of *Max* in the T cell lineage using lck-cre led to marginally impaired differentiation of double negative (DN) to double positive thymocytes (DP) (Supplemental Fig. S1G), Taken together, our data indicate that *Max* loss attenuates overall lymphocyte development, rather than completely abolishing it.

### **Requirement for *Max* in Activated lymphocytes and Eµ-*Myc*-induced lymphomagenesis**

To study the requirement for *Max* in situations where *Myc* expression is elevated, we activated B220+ B-cells *in vitro* using bacterial lipopolysaccharides (LPS). We found activation to be severely compromised in *Max* KO mice, and B-cells exhibited little increase in cell size (Fig 2A). This was accompanied by reduced cell numbers, viability and apoptosis compared to LPS-treated controls (Fig. 2B-D, Supplemental Fig. S2E). *Max* KO B-cells also failed to proliferate when activated with IgM-µ or a combination of anti-CD40/ IL-4 *e*x *vivo* (Supplemental Fig. S2A-E). Similar effects on proliferation and cell size were observed in *Max*^fl/fl^ lck-Cre CD3+ T lymphocytes stimulated with anti-CD3 and anti-CD28 (Supplemental Fig. S2F, G).

**Fig. 2:**
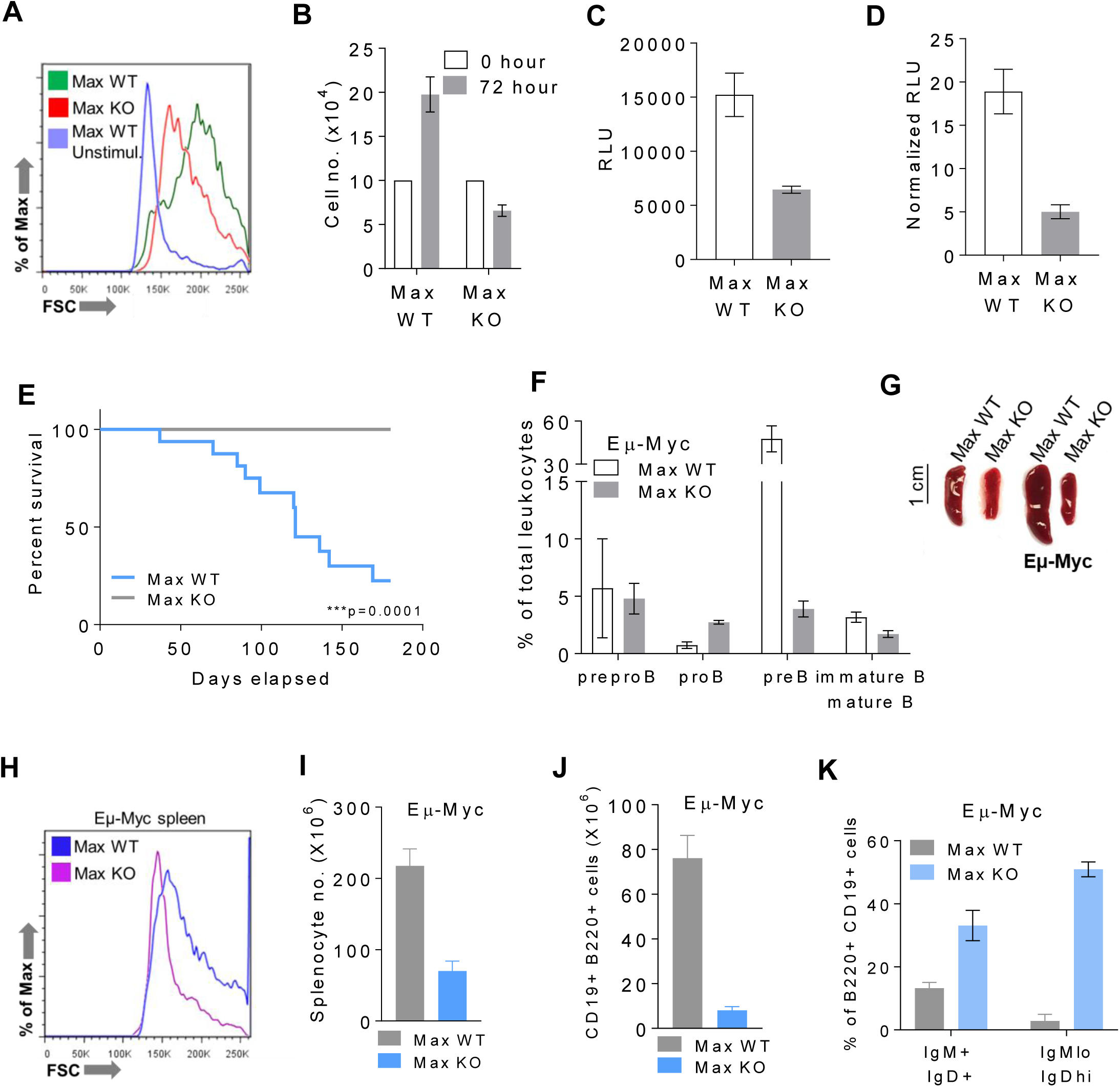
Requirement for *Max* in Activated B-cells and Eµ-*Myc*-induced lymphomagenesis. (A) Cell size as determined by forward scatter in WT unstimulated and LPS activated *Max* WT and *Max* KO B220+ cells. (B) Cell number 72 hours post treatment in LPS treated *Max* WT and KO (n=10 from 5 WT and KO mice). (C) cell viability and (D) apoptosis in LPS activated B-cells (n=9 from 3 WT and KO) assessed using luciferase-based cell titer glo and caspase glo assays (E) Kaplan Meier curve showing survival analysis of Max WT (n=23) and KO (n=14) Eµ-Myc animals up to 180 days. P-value calculated using Log-rank (Mantel-Cox) test. (F) Analysis of B-cell precursor populations in Eµ-*Myc* BM. (G) Spleens from normal and Eµ-*Myc* mice. (H) Histogram of cell size of Eµ-*Myc Max* WT and KO mice. Splenocyte number (I) and CD19+ B220+ cells (J) in Eµ-*Myc* WT and KO mice (n=3 for each). (K) Proportion on mature IgD positive B-cells in Eµ-*Myc* spleens (n=3).

To ascertain whether *Max* loss affects MYC-driven lymphomagenesis, we crossed our *Max* conditional allele with Eµ-*Myc* mice in which the *Myc* transgene is predominantly restricted to the B lymphoid lineage (Adams et al. 1985; Harris et al. 1988). While all the Eµ-*Myc Max* WT animals developed B-cell lymphomas with a median survival of 97 days, none of the Eµ-Myc Max KO mice developed lymphoma even out to 300 days (Fig. 2E, data not shown). Pre-malignant Eµ-*Myc* BM B-cell precursors exhibited developmental defects, including a block at the pre-B-cell stage (Langdon et al. 1986). Analysis of BM populations failed to show an expansion of a pre-B-cell population in Eµ-*Myc Max* KO mice (Fig. 2F). In addition, B220+ cells from Eµ-*Myc Max* KO mice were smaller than Eµ-*Myc* controls and exhibited decreased total RNA content (Supplemental Fig. S2H, I). Augmented spleen size and splenocyte numbers are typical of Eµ-*Myc* induced B-cell lymphomagenesis (Harris et al., 1988). Compared with Eµ-*Myc* Max KO mice, Eµ-*Myc* Max WT mice exhibited increased spleen size, and significantly increased numbers and cell size of total and B220+ CD19+ splenocytes (Fig 2. G-J). Eµ-*Myc Max* KO B220+ splenocytes also had an increased proportion of mature IgM and IgD positive B-cells when compared to wildtype controls (Fig. 2K) indicating the *Max* KO cells do not exhibit the defects in differentiation characteristic of pre-malignant Eµ-*Myc* cells. These data demonstrate that Eµ-*Myc* lymphomagenesis is severely compromised in the absence of *Max.*

### ***Max* loss affects E2F targets and pro-inflammatory pathways in B-cells**

To determine the effects of *Max* loss on the transcriptional program of normal and Eµ-*Myc* expressing B-cells we performed RNA-seq using B220+ cells of the four genotypes described above. Strikingly, *Max* deletion in normal B-cells doesn’t completely phenocopy *Myc* loss in B-cells. First, MAX depletion does not perturb the expression of B-cell lineage transcription factors (Supplemental Fig. S3A), in contrast to MYC loss, which was previously shown to downregulate expression of factors such as EBF and PAX5 (Vallespinós et al. 2011). Second, *Max* KO B-cells exhibit a significant up-regulation of genes involved in pro-inflammatory pathways (Fig. 3A, Supplemental Fig. S3B, C). *Max* KO B220+ cells were also consistently larger than controls (Supplemental Fig. S3D). Taken together, our data suggest that these cells are in a quasi-activated state, possibly related to the loss of repressive MAX dimers (e.g. MNT-MAX see below), compensating for the absence of MYC-MAX function. Interestingly, a majority of genes that exhibit decreased expression upon *Max* knockout are cell cycle related (Fig. 3B) and include E2F targets. Consistent with the loss of GC cells (Fig.1H, I), a group of genes crucial for GC maintenance are also down-regulated in *Max* KO B-cells (Supplemental Fig. S3E).

**Fig. 3:**
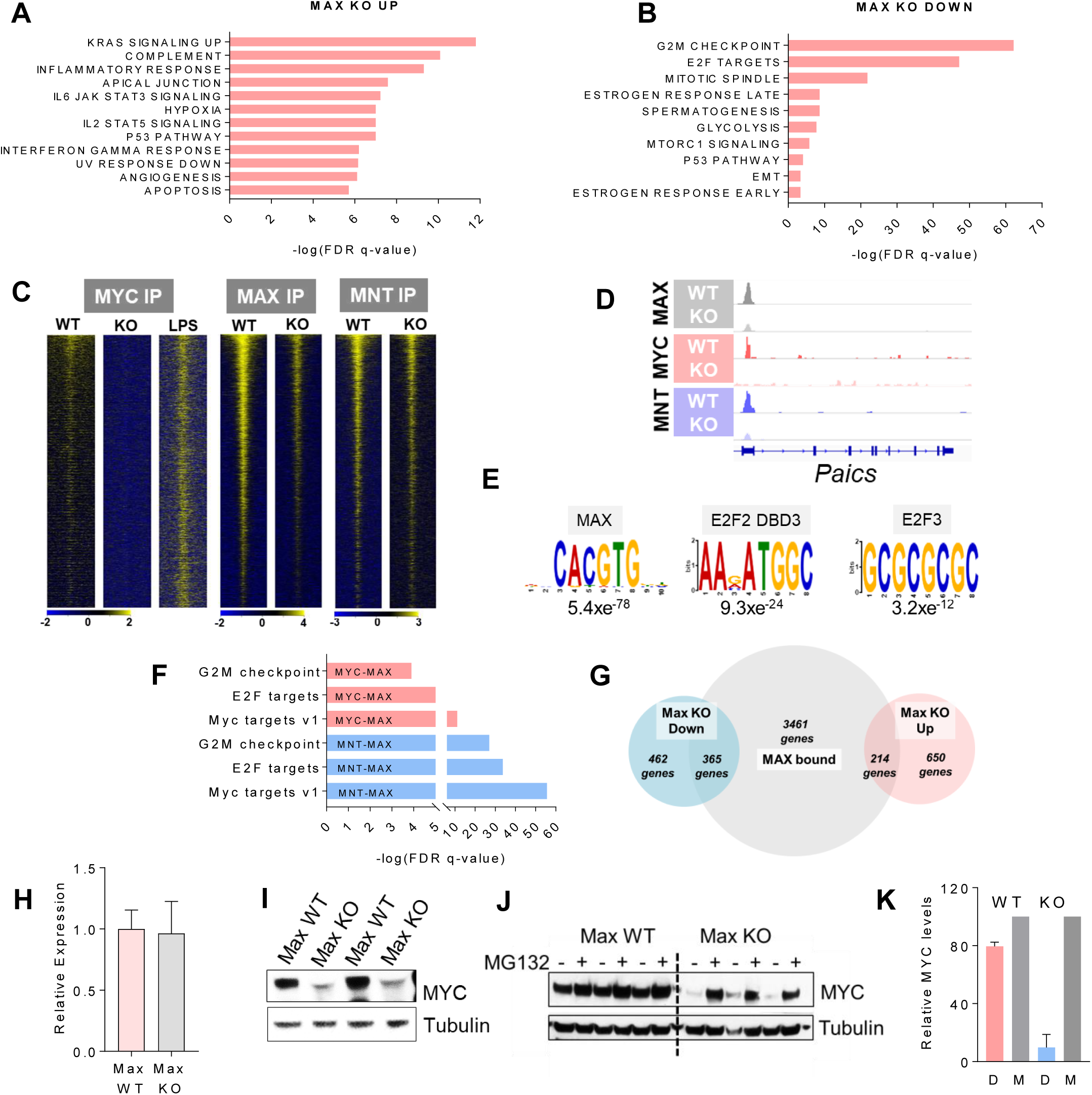
Gene expression profiling and genomic occupancy of Max in B-cells. Hallmark gene set enrichment for pathways (A) upregulated and (B) downregulated in Max KO B-cells relative to WT B-cells. (C) NGS plots depicting genomic occupancy of MYC, MAX and MNT in Max WT and KO B-cells ranked on expression changes. X-axis showing (D) Representative peaks for MAX, MYC and MNT at E2F target *Paics*. (E) Motifs significantly enriched at MAX bound genes. (F) Gene set enrichment for pathways enriched in MAX bound genes. (G) Overlap of MAX bound genes with genes that are differentially expressed in *Max* KO B-cells (FDR<0.05 cutoff for differential expression). (H) mRNA and (I) protein levels of *Myc* in WT and KO B-cells (n=4 for WT and KO, error bars represent SEM. (J) Immunoblot and (K) quantification of MYC levels following 2hours MG132 treatment of *Max* WT and KO B-cells

To identify direct MYC-MAX and MXD-MAX targets, we carried out genomic occupancy analysis using CUT&RUN on *Max* WT and KO B-cells. CUT&RUN (cleavage under targets and release using nuclease) sequencing utilizes a combination of antibody targeted controlled cleavage and nuclease-based release of DNA fragments to analyze protein occupancy on DNA (Skene and Henikoff, 2017; Janssens et al. 2018). MAX was found to bind to 4040 gene loci (within 10 kb of the TSS) in WT B-cells and there was an overall reduction in MAX, MNT and MYC binding in Max KO cells when compared to WT (Fig. 3C). MYC occupancy was low in WT when compared to LPS stimulated B-cells and decreased further in MAX null B-cells. The decrease in MAX binding at a representative gene is shown in Fig. 3D. The MAX binding observed in the *Max* KO B-cells was most likely derived from a fraction of B-cells that escaped Cre-mediated deletion of *Max* (∼14%, see Fig. 1E). In addition to the E-Box motif, MEME and HOMER analysis revealed a significant enrichment for E2F motifs in MAX bound regions in WT B-cells compared to the IgG control (Fig. 3E, Supplemental Fig S3F). We observed a substantial overlap between MAX, MYC and MNT binding in WT B cells (Supplemental Fig S3G). GSEA revealed that cell cycle and E2F target gene sets were enriched in the MNT-MAX and MYC-MAX bound gene populations whereas there was no significant enrichment for inflammation related genes (Fig 3F). Therefore, the effects of *Max* inactivation on upregulation of inflammatory genes (Fig 3A, Supplemental Fig S3B, C) are likely to be indirect.

When we correlated MAX occupancy with gene expression changes in *Max* null cells, we found that approximately half the genes downregulated in *Max* null cells were occupied by MAX in WT cells, while 25% of upregulated genes were directly bound by MAX (Fig. 3G). Remarkably, 86% of MAX-bound genes were not differentially expressed in *Max* KO B-cells including the majority of MAX bound E2F target genes. Expression of E2F1-3 and phosphorylation of Rb were also unaffected by *Max* loss (Supplemental Fig S3H, J). The lack of change in expression of a majority of MAX bound genes may be due to loss of binding by both the transcriptionally activating (MYC) and repressive (MXD, MNT, MGA) heterodimerization partners of MAX and is consistent with the weak effects of MAX deletion on B-cell differentiation. Consistent with this idea, we observed that the expression of MYC and E2F targets genes that are bound by MYC, MNT and MAX remained unchanged upon the deletion of Max (Supplemental Fig S3I).

To determine whether the gene expression changes and decreased MYC occupancy in *Max* KO cells is solely due to the inability of MYC to bind DNA without MAX, we measured MYC levels in *Max* KO B-cells. While *Myc* mRNA levels where not affected by *Max* deletion (Fig. 3H) we observed a striking reduction in MYC protein levels in Max KO cells (Fig 3I, Supplemental Fig S3K). This result was surprising in light of previous studies indicating that MYC is subject to negative auto-regulation in normal B-cells (Grignani et al, 1990). Importantly, however, we found that treatment of B-cells with the proteasomal inhibitor MG132 resulted in near complete restoration of MYC levels in *Max* KO B-cells (Fig 3J). Taken together, these results strongly suggest that MAX influences MYC stability. Phosphorylation of AKT (p-AKT) and subsequent phosphorylation of GSK3β at Ser9 is known to increase MYC stability (Cross et al, 1995; Farrell and Sears, 2014). Because levels of both p-AKT Ser473 and p-GSK3β (Ser9) appeared to be higher in Max KO B-cells (Supplemental Fig. S3J, L), we surmise that MAX loss regulates factors independent of GSK3β-mediated regulation of MYC stability.

### ***Max* loss leads to a global down-regulation of the MYC signature in Eµ-*Myc* expressing B-cells**

When we examined whether *Max* deletion in premalignant Eµ-*Myc* cells impacts MYC levels we observed a striking decrease in MYC protein levels and half-life (Fig. 4A, Supplemental Fig. S4A, B,C) although mRNA levels were only modestly reduced (Fig. 4B, Supplemental Fig. S4D). In this scenario, we hypothesized that the loss of MYC stability would have a widespread effect on the transcriptional profile of *Max* null cells. Indeed, in contrast to normal B-cells, where the loss of *M*ax affects the expression of ∼550 genes (2 fold change cutoff, FDR < 0.05), MAX depletion in Eµ-*Myc* mice causes a dramatic shift in the expression of thousands of genes (Fig. 4C), with Eµ-*Myc Max* WT cells occupying a space distinct from the other genotypes in a Principle Component Analysis plot (Fig. 4D). Whereas genes upregulated in Eµ-*Myc Max* KO cells showed a close alignment with signatures enriched in normal B-cells lacking *Max* (Fig 4E), genes downregulated in Eµ-*Myc Max* KO cells revealed a significant enrichment for MYC signatures (Fig 4F). This translated to robust differences in expression where Eµ-*Myc Max* KO and *Max* WT profiles closely resemble each other and are nearly the inverse of the Eµ-*Myc Max* WT (Fig. 4G). Interrogation of a publicly available ChIP-seq dataset in B-cells (Sabo et al. 2014) revealed that 73% of the genes known to be bound by MYC in Eµ-*Myc* pre-malignant cells are differentially expressed in Eµ-*Myc Max* KO B220+ cells (Fig. 4H, I, Supplemental Fig S4E). We also observed that in contrast to normal B-cells, a substantial proportion of MAX regulated inflammation related genes are directly bound by MYC (Fig. 4J). In addition, a large fraction of MYC bound E2F targets (Kuleshov et al. 2016) (Fig. 4K) are actually down-regulated in Eu-MYC MAX KO cells. This may be due partly to the decrease in E2F1-3 expression in KO cells (Supplemental Fig. S4F).

**Fig. 4:**
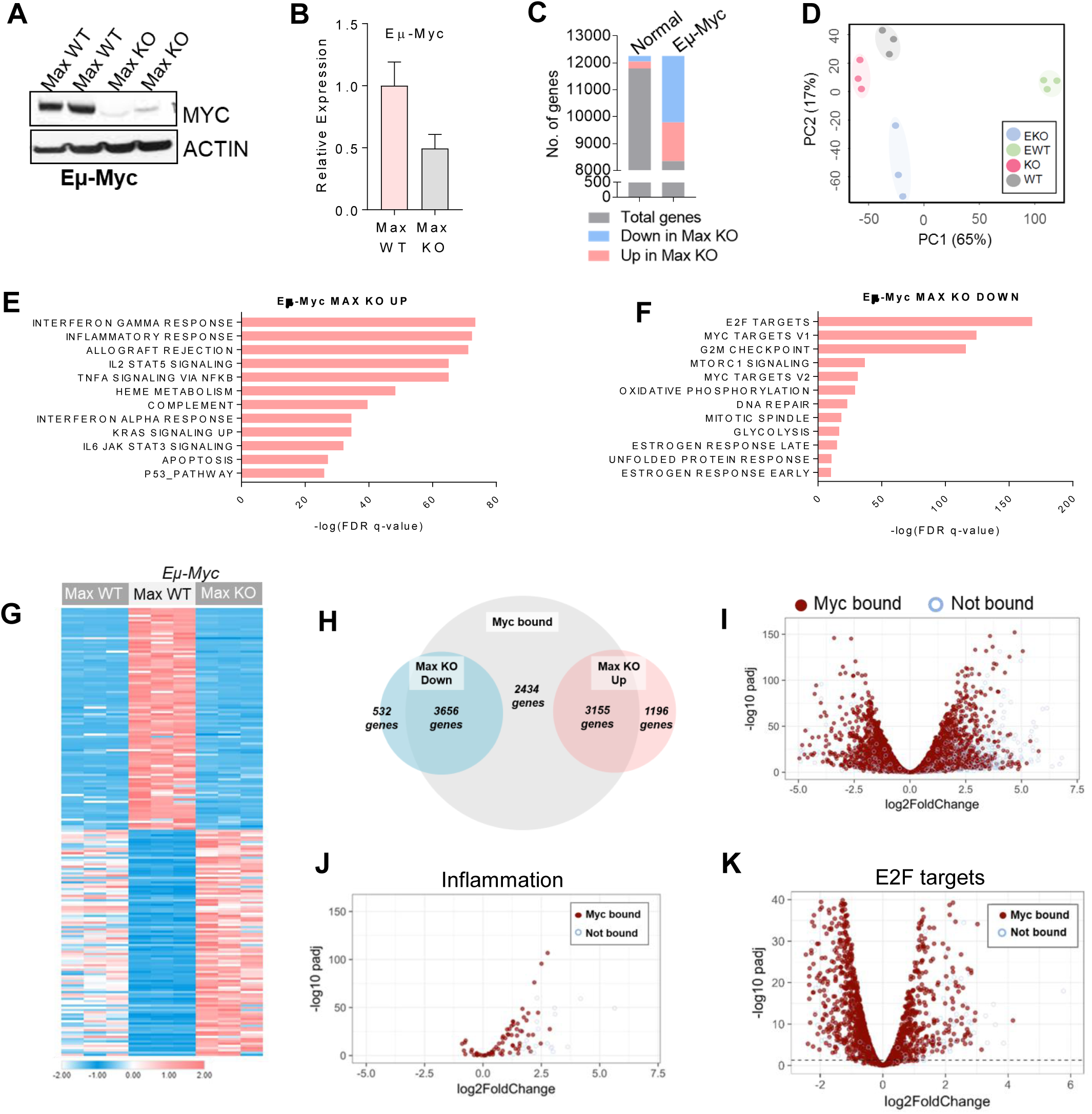
*Max* loss leads to a global down-regulation of the Myc signature in Eu-*Myc* premaligant cells. (A) MYC protein and (B) *Myc* mRNA levels in Eu-Myc Max WT and KO cells. (C) Summary of total differentially expressed genes in *Max* KO normal and pre-malignant B-cells. (D) Principal component analysis of all four genotypes (EWT= Eu-*Myc Max* WT, EKO = Eu-*Myc Max* KO, WT = *Max* WT and KO= *Max* KO). (E, F) Hallmark gene set enrichment for pathways upregulated (E) and downregulated (F) in Eu-*Myc* pre-malignant *Max* KO B-cells. (G) Heatmap representation of global transcriptional changes in Eu-*Myc* pre-malignant *Max* WT and *Max* KO cells. (H) Venn diagram and (I) Volcano plot of differentially expressed genes that are directly bound by MYC. (J, K) Volcano plots depicting proportion of differentially expressed (J) inflammation related genes and (K) E2F target genes that are directly bound by MYC in Eu-*Myc* pre-malignant cells.

Together, our data reveal that *Max* loss in the context of Eµ-*Myc* leads to an unstable pool of MYC and largely reverses the transcriptional effects of MYC. Our results indicate that MYC-MAX cooperation with E2F factors is required to drive proliferation in Eµ-MYC pre-malignant cells. Our findings also suggest that MYC-MAX genomic binding and transcriptional activity is not absolutely required for several key aspects of early B-cell differentiation. Nonetheless, deletion of *Max* attenuates certain processes where MYC is critically required, such as GC formation. However, in situations where MYC is overexpressed and dysregulated, such as in the Eµ-*Myc* model, cells become hyper-dependent on MYC. Here, Max loss and subsequent MYC destabilization leads to the global downregulation of thousands of MYC targets with dramatic phenotypic consequences.

### MAX loss or inhibition of MYC-MAX dimerization results in repression of MYC stability factors

Given the profound effect of MAX loss on MYC stability, we wanted to identify effectors downstream of MYC-MAX that may form a positive feedback loop to maintain MYC stability. While MYC proteins generally have short half-lives, on the order of 20-30 minutes, MYC half-life increases in several Burkitt’s lymphoma lines and ES cells (Gregory and Hann, 2002, Hann and Eisenman 1984, Cartwright et al, 2005). The rate of MYC protein degradation is mediated by several factors that interfere with signals triggering MYC ubiquitination and proteasomal degradation (Farrell and Sears, 2014). We noted in our RNA-seq data that genes encoding MYC stability factors, including *Btrc, Cip2a* and *Set,* are upregulated in Eµ-*Myc* cells, relative to normal B220+ cells, and downregulated in Eµ-*Myc Max* KO B-cells (Fig 5A). Moreover, the promoters of these genes are directly bound by MYC in premalignant cells (Fig. 5B, Supplemental Fig S5A). CIP2A and SET are inhibitors of protein phosphatase 2A (PP2A) which normally dephosphorylates the stabilizing phospho-serine 62 (S62) within the conserved Myc Box I (MB1) phospho-degron. CIP2A and SET are often overexpressed in tumors and block PP2A activity, resulting in persistence of phospho-S62 and MYC (Junttila and Westermarck, 2008, Junttila et al. 2007). BTRC functions to enhance MYC stability via ubiquitylation (Popov et al. 2010).

**Fig. 5:**
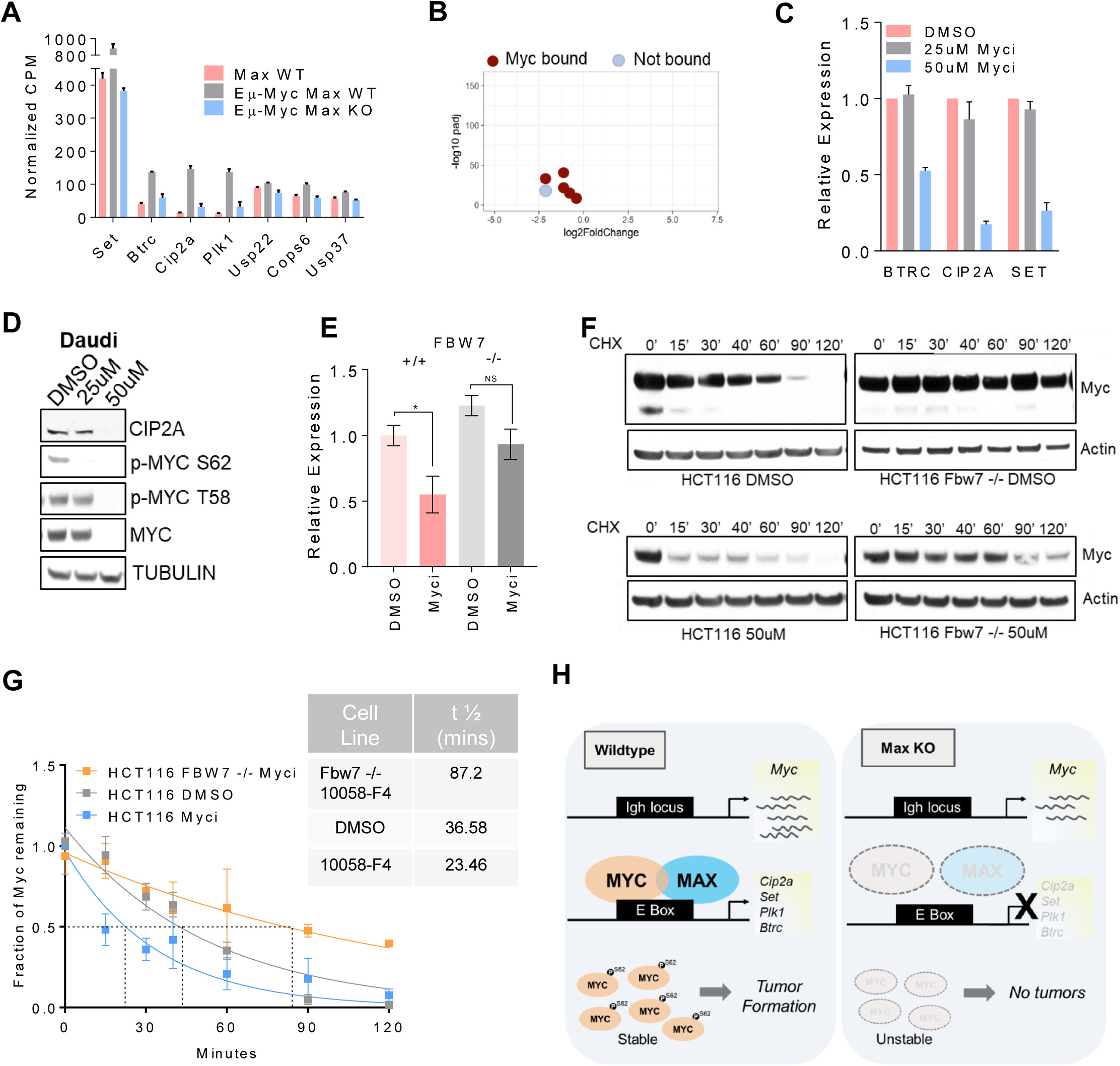
Factors mediating MYC degradation in the absence of MAX. (A) Normalized expression values of MYC stabilizing genes in WT and Eµ-*Myc* B220+ cells. (B) Volcano plot showing expression changes for MYC stability genes and ChIP binding data for MYC in pre-malignant Eµ-*Myc* B220+. (C) qPCR for *Btrc Cip2a* and *Set* in DMSO and Myci treated Daudi cells. (D) Western blot for pMYC S62 and CIP2A levels in Myci treated Daudi cells. (E) qPCR for CIP2A levels in HCT116 cells (n=3, p=0.048 for Myci vs control in WT). (F) Representative blot for MYC levels in HCT116 and HCT116 FBW7 -/- cells following a cycloheximide chase. (G) Determination of MYC half-life in Myci treated control and FBW7 -/- cells (n=3 experiments). (H) Model depicting proposed auto-regulation of MYC by MAX *in vivo* in pre-malignant Eµ-Myc cells.

To confirm whether these genes are regulated by MYC-MAX in tumor lines, we treated Daudi and P493 human B-cell lymphoma lines (Schuhmacher et al. 2001) with 10058-F4 (referred to herein as Myci), a small molecule reported to inhibit MYC-MAX heterodimerization (Yin et al. 2003). We observed a decrease in MYC protein levels, accompanied by reduced proliferation and a downregulation of the stability factors in Myci treated cells (Fig. 5C, Supplemental Fig. S5 B-E). While *Myc* RNA levels were only moderately reduced (Supplemental Fig. S5F) MYC phospho-serine62 levels were nearly eliminated upon Myci treatment (Fig. 5D). In addition, a time-course of MYC degradation following cycloheximide treatment in P493-6 cells revealed that MYC half-life is reduced in Myci treated cells (Supplemental Fig. S5G, H). Similar results were obtained in colon, pancreatic and lung adenocarcinoma human tumor lines (Supplemental Fig. S6A-F). ENCODE data shows that MAX is directly associated with the promoter-proximal regions of the *Btrc, Cip2a* and *Set* genomic loci in HCT116 cells (Supplemental Fig. S6G). To confirm that the decrease in MYC protein levels is indeed through the loss of multiple stability factors that affect *Fbw7* activity, we examined MYC levels in HCT116 colon cancer cells lacking the FBW7 ubiquitin ligase known to target MYC for degradation (Welcker et al. 2004; Yada et al. 2004). We found that MYC turnover in FBW7-/- HCT116 is largely unaffected by treatment with the dimerization inhibitor compared to the rapid turnover in control HCT116 cells treated with inhibitor (Fig. 5F, G). Moreover, CIP2A levels are unchanged in FBW7 null cells upon Myci treatment whereas they decrease in Myci treated WT HCT116 cells (Fig 5E). This supports the notion that disruption of MYC-MAX dimerization doesn’t affect MYC-MAX mediated transcription alone. The augmented degradation of MYC in cells where MYC-MAX dimerization is disrupted contributes to the decrease in MYC target transcription. Our data suggest that the decreased stability of MYC is at least partially dependent on the activity of the FBW7-SCF ubiquitination complex.

Lastly, we asked whether the MYC paralogs N-MYC and L-MYC are also destabilized by loss of heterodimerization. A previous study reported a decrease in N-MYC in SKNBE neuroblastoma cells treated with 10058-F4 (Zirath et al. 2013). We extended our analysis to another *Mycn* amplified neuroblastoma line IMR-32 and the *Mycl* amplified small cell lung cancer line H-2141. Treatment with Myci leads to a reduced growth and a decrease in N- and L-MYC protein respectively (Supplemental Fig. S7A, B; Supplemental Fig. S7C, D).

Our findings suggest that specific MYC-MAX dimerization inhibitors will be doubly efficacious, targeting both MYC driven transcription and MYC protein levels. In addition, by eliminating MYC, dimerization inhibitors would suppress transcription independent functions of MYC, such as those mediated by MYC-nick (Conacci-Sorrell et al. 2010). This additional layer of autoregulation of MYC stability is likely to have broad mechanistic consequences during tumor initiation. Our data lends support to the notion that transcriptional upregulation of PP2A inhibitors by MYC may lead to enhanced MYC stability and function in pre-malignant settings to facilitate transformation, even in the absence of genomic alterations of *Myc* (Junttila and Westermarck, 2008). It will therefore be important to closely examine the consequences of MYC inhibition in cancers where PP2A is lost and/or regulators like CIP2A and SET are over-expressed.

In summary, our data suggest that loss of *Max in vivo* disables MYC activity through inhibition of direct MYC association with genomic DNA and through destabilization of the MYC protein itself. We surmise that increased MYC degradation is facilitated by decreased MYC-MAX activity at the promoters of genes such as *Cip2a*, whose protein products normally serve to stabilize MYC by attenuating FBW7 mediated proteasomal degradation (Fig 5H) (Junttila et al. 2007). In this scenario MYC-MAX heterodimers drive a feed-forward circuit that reinforces high MYC expression in tumors as diverse as B-cell lymphomas and colon adenocarcinomas. In fact, a recent study shows that MYC regulates the kinase *Plk1* to maintain its stability in an aggressive form of lymphoma (Ren et al. 2018). Disruption of this circuit, for example, in Eµ-*Myc, Max* null B-cells results in loss of MYC protein expression and a complete abrogation of lymphomagenesis. Surprisingly, the major hallmarks of normal B-cell differentiation are largely unperturbed by *Max* loss. The fact that MYC protein is destabilized and nearly absent in the *Max* null B-cells makes it unlikely that MYC is functioning independently of MAX. This raises the possibility that the requirement for MYC activity is rather minimal, or readily compensated for by factors such as E2Fs in normal B-cell progenitors. In *Max* KO cells, loss of repressive MAX-MXD heterodimers may serve to partially compensate for diminished MYC activity. For example, MNT binds thousands of promoters in normal B-cells and its binding is partially attenuated in MAX KO cells (Fig 3C). This alleviation of repression may also underlie the indirect activation of pro-inflammatory pathways in *Max* null B-cells. Nonetheless, normal B-cell development overall is impaired by B-cell specific deletion of *Max* as evidenced by markedly decreased B-cell population sizes in both bone marrow and spleen, with a nearly complete loss of germinal center cells in the latter. Our data underscore the complex circuitry, and the integrated context-dependent functions of the MAX and the broader MYC network.

## MATERIALS AND METHODS

All mice were housed and treated according to the guidelines provided by the Fred Hutch Institutional Animal Care and Use Committee. For *Max* conditional mice, clonal G418R-targeted ES cell lines were produced, and integration of the targeting vector verified by genomic PCR and Southern blotting. For B-cell developmental studies, cells harvested from bone marrow and spleen were stained with fluorochrome conjugated B-cell lineage specific markers. They were then analyzed on a BD FACS Canto II. Immunohistochemistry on mouse spleens was performed after heat-induced antigen retrieval. B-cells were sorted from bone marrow using an AutoMACS mouse B-cell isolation kit and stimulated *ex vivo* with LPS. Cell titer glo and caspase glo luciferase assays (Promega) were utilized to assess cell growth and apoptosis. All RNA isolation was performed using the Direct-zol RNA Miniprep kit (Zymoresearch). For RNA-seq B220+ cells were purified using B220+ microbeads (AutoMACS) from spleens of 5-6-week-old mice from all genotypes, RNA integrity assessed using Tapestation (Agilent) and libraries prepared using the TruSeq RNA Sample Prep v2 kit with 500ng input RNA. Paired-end sequencing was performed on an Illumina HiSeq 2500. Reads were aligned to mm10 mouse reference genome using STAR2 and normalized using EdgeR. Gene ontology analysis was performed using hallmark datasets on mSigDB (Subramanian et al, 2005; Liberzon et al, 2015). Heatmaps were generated using Morpheus (https://software.broadinstitute.org/morpheus). For chromatin occupancy studies, primary mouse B-cells isolated from spleens were prepared fresh or stimulated ex vivo with LPS. Cells were bound to ConA beads, then permeabilized, and incubated overnight with antibody to MYC, MAX and MNT. One million cells were used per IP and the Cleavage Under Targets and Release Using Nuclease (CUT&RUN) protocol was followed (Janssens et al, 2018). 25×25 paired-end sequencing was performed (5-10 million reads) on an Illumina HiSeq 2500 instrument and sequences were aligned to the mm10 reference genome assembly using Bowtie2. Peak calling employed a threshold peak calling script to differentiate signal to noise. This processing was carried out with bedtools, custom R scripts defining genome position, and the GenomicRanges R package. For MYC peak calling in WT B cells, only peaks that were identified in both WT B cells and LPS stimulated B cells were considered real to mitigate background issues. De novo enrichment for sequence specificity was determined using Homer (Heinz et al, 2010) and MEME-ChIP (Machanick and Bailey, 2011). Genomic plots were made using ngs.plot (Shen et al, 2014) or the R package ggplot2. For qRT-PCR, 500ng-2μg of input total RNA was used and cDNA generated using the Revertaid cDNA synthesis kit (Thermo-Fisher). SYBR green and FAM based methods were used to perform qPCR on a Biorad iCycler. For western blots, cells were lysed in RIPA buffer with phosphatase and protease inhibitors. For inhibitor studies, cells were treated with the MYC inhibitor 10058-F4 for 2 days prior to harvest. Cells were treated with 5μg/μl Cycloheximide and 10μM MG132. Daudi cells were grown in RPMI with 10% FBS and HCT116 lines were maintained in DMEM with 10% FBS. Refer to Supplemental Methods for lists of reagents used.

## Supporting information

Supplemental Data: Max deletion destabilizes MYC protein and abrogates Eu-Myc lymphomagenesis

## ACKNOWLEDGEMENTS

We thank Nayanga Thirimanne, Bruce Clurman, Arnaud Augert, David MacPherson, Nan Hyung Hong and Patrick Carroll for providing essential reagents. We also acknowledge Genomics, Scientific Imaging, Flow Cytometry and Experimental Histopathology scientific resources at Fred Hutch. This work was supported by NIH/NCI grants RO1 CA057138 and R35 CA 231989 (to R.N.E).

