## Supplemental Data: Max deletion destabilizes MYC protein and abrogates Eu-Myc lymphomagenesis for "*Max* deletion destabilizes MYC protein and abrogates Eμ-*Myc* lymphomagenesis"

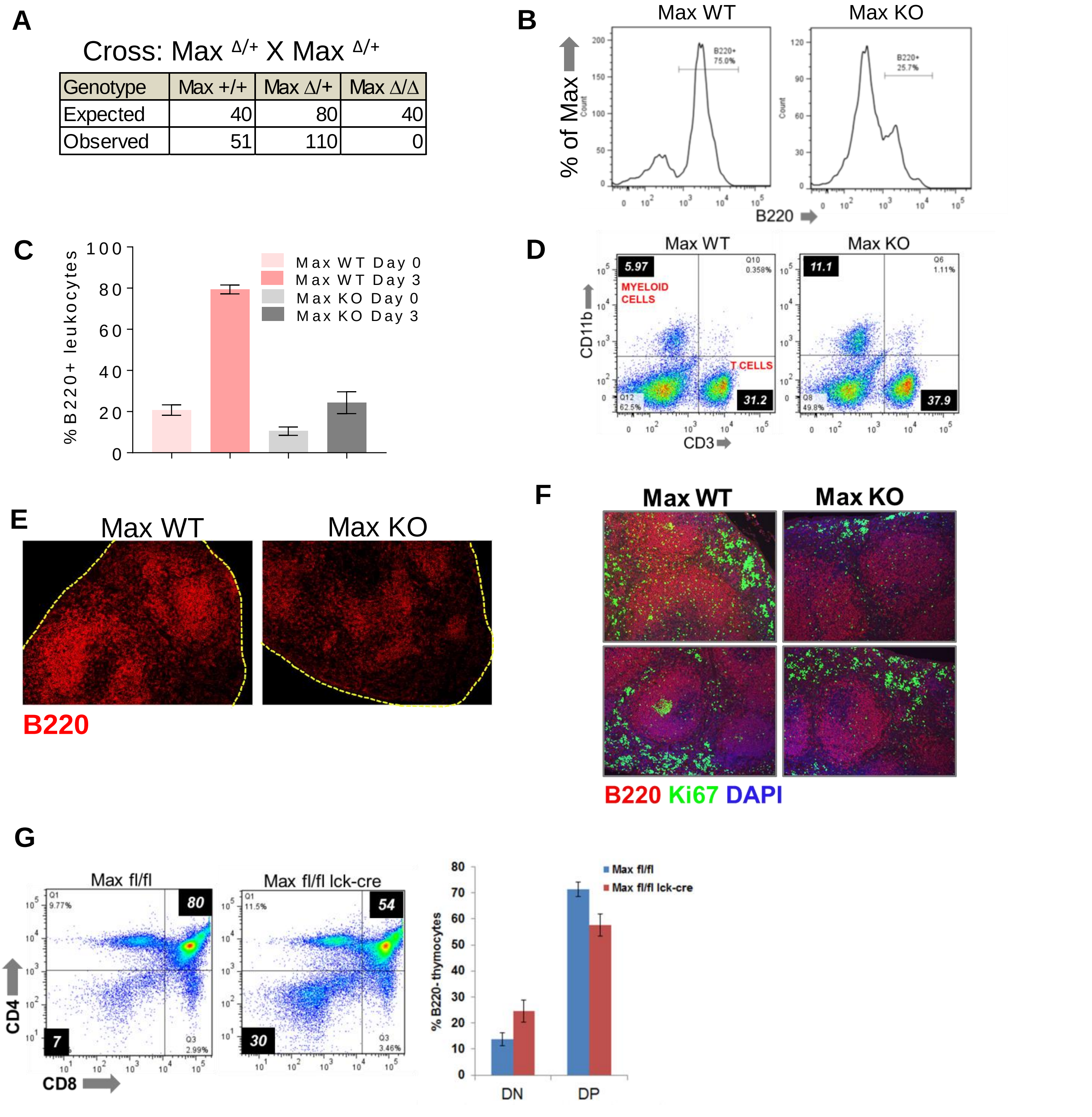


**Supplemental Fig. S1 Deletion of Max impairs B-cell Development.**

(A) Expected and observed frequencies from *Max* heterozygous crosses. (B) Representative histogram and (C) quantification of B220+ cell percentage 72 hours post stimulation of BM *ex vivo* with Il-7. (D) Proportion of Myeloid (CD11b+) and T-cell (CD3+) in *Max* WT and KO spleens. (E) Representative staining for B220 on WT and KO spleens. (F) Double IF for B220 and Ki67 in *Max* WT and KO spleens. (G) Proportion of double negative (DN) and double positive (DP) thymocytes in *Max*$^{fl/fl}$ and *Max*$^{flfl}$ lck-cre mice.

### Supplemental Fig. S2

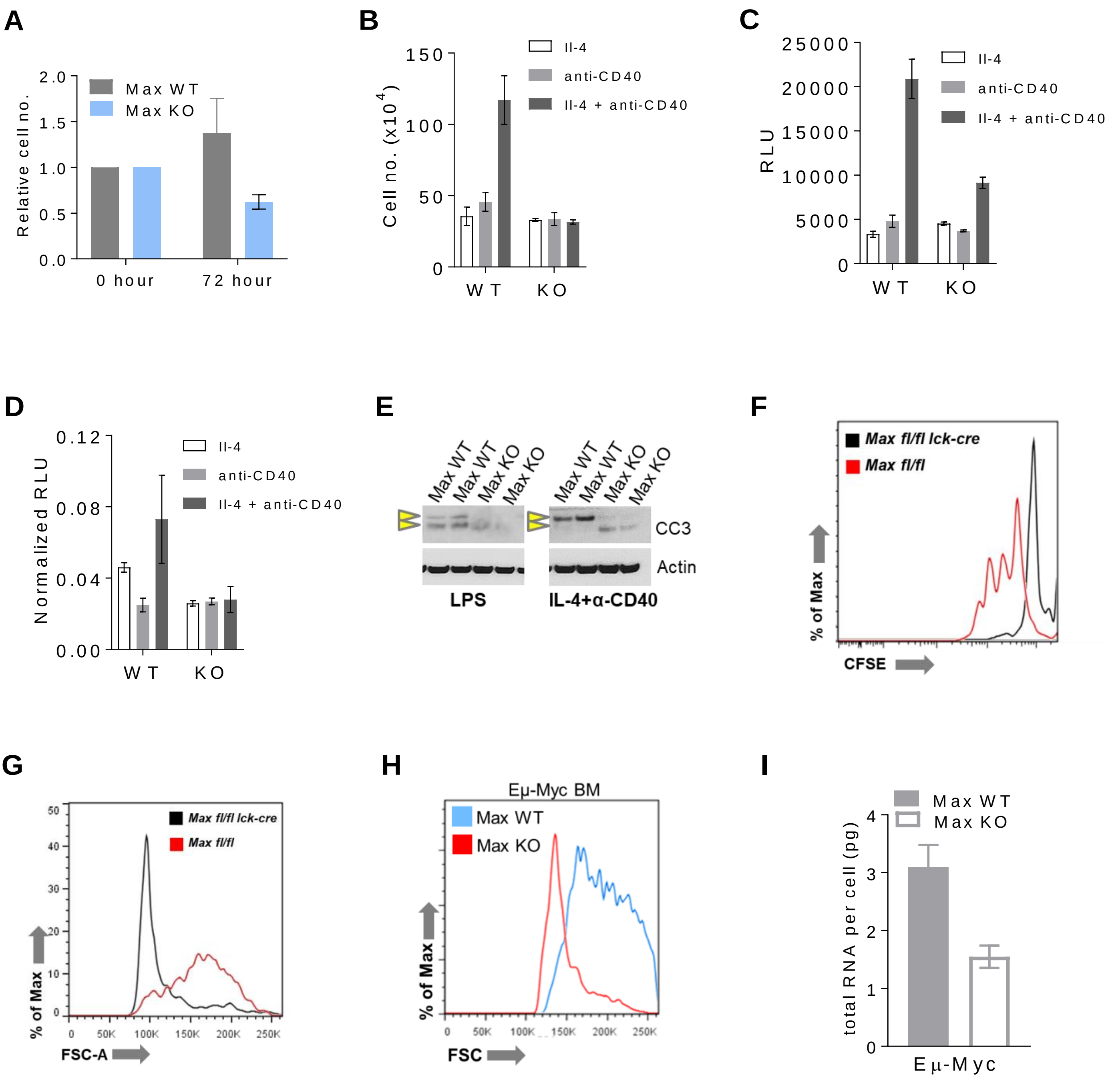

**Supplemental Fig. S2 Requirement for *Max* in Activated B-cells and Eμ-*Myc*-induced lymphomagenesis.** (A) Relative cell number in IgM μ activated B220+ cells 72 hours post treatment (n=2 WT n=3 KO). (B) Relative cell number, (C) relative viability assessed using cell titer glo (D) normalized caspase activity in Max WT and KO cells 72 hours post treatment with Il-4, anti-CD-40 or a combination of both (n=6 from 2 WT and KO mice). (E) Cleaved caspase 3 levels in stimulated vs. non-stimulated WT and KO B-cells. (F) Representative histogram of CFSE dilution following activation of CD3+ T cells from control and lck-cre mice 72 hours post treatment with anti-CD3/anti-CD28. (G) Cell size of anti-CD3 anti-CD28 activated T cells from control and lck-cre animals. (H) Size of B220+ cells isolated from the spleens of pre-malignant Eμ-*Myc* *Max* WT and KO mice. (I) Total RNA content per cell of Eμ-*Myc* *Max* WT and *Max* KO B220+ cells.

### Supplemental Fig. S3

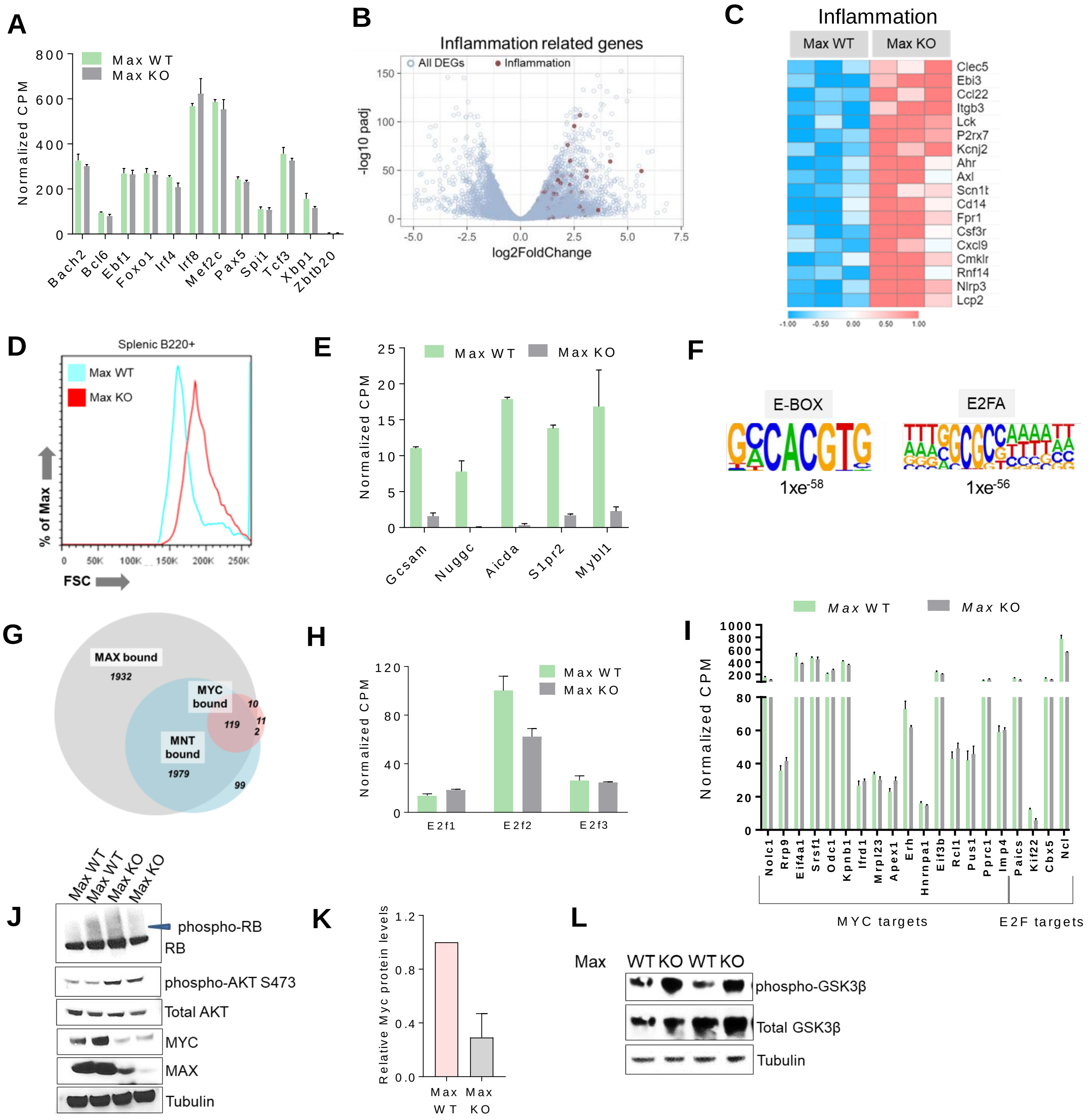

**Supplemental Fig. S3 Gene expression profiling and genomic occupancy of MAX in B-cells**

(A) Normalized CPM values for genes that encode transcription factors critical for B-cell development comparing *Max* WT and KO B-cells. (B) Volcano plot and (C) heatmap showing a general upregulation of inflammation related transcripts in *Max* KO B-cells. (D) Representative histogram depicting difference in cell size between *Max* WT and KO B-cells. (E) Normalized CPM values for genes that are important for germinal center function (F) HOMER analysis for E2F and E-Box motifs on MAX occupied promoters in WT cells. (G) Venn diagram showing overlap between MYC, MNT and MAX occupancy in WT B cells. Numbers denote genes (H) Expression of E2F1-3 in *Max* WT vs. KO cells. (I) Normalized CPM values of MNT, MAX and MYC bound MYC and E2F targets in WT and KO B cells. (J) RB and p-Akt Ser 473 levels in Max WT and KO cells (K) Densitometry of MYC protein levels in *Max* WT and KO cells (n=4 for each) (L)phospho-GSK3b Ser 9 levels in *Max* WT and KO B-cells (same immunoblot as Fig. 3H).

### Supplemental Fig. S4

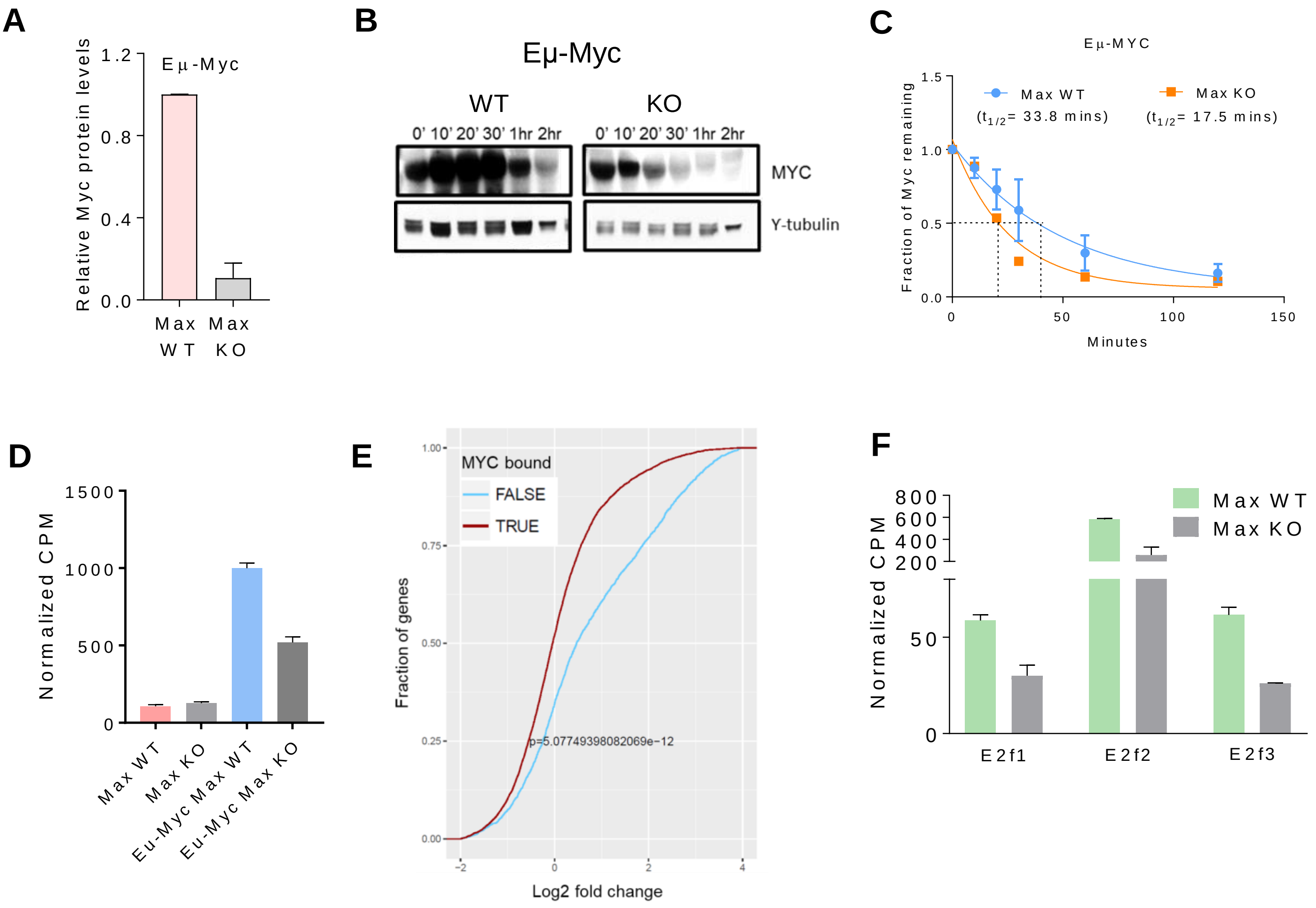


**Supplemental Fig. S4 *Max* loss leads to a global down-regulation of the MYC signature in Eμ-*Myc* premaligant cells.** (A) densitometry for MYC levels in WT and KO cells. (B) Immunoblot for MYC and (C) determination of half-life using CHX chase on a pool of Eμ-*Myc Max* KO and Eμ-*Myc Max* WT B220+ cells (n=3). (D) normalized CPM values from RNA-Seq data for *Myc* expression in pre-malignant Eμ-*Myc* B220+ cells (E) Cumulative distribution plot showing a significant enrichment for genes that are differentially expressed in Eμ-*Myc Max* KO B220+ cells and are also directly bound by MYC (F) Normalized expression of E2F family members in Eμ-*Myc Max* WT vs. KO cells (FDR<0.05). n=3 for WT and KO unless otherwise noted.

### Supplemental Fig. S5

**A**

| Gene | LFC | FDR | Myc binding | Pol2 binding |
|---|---|---|---|---|
| *Cip2a* | -2.2 | 4.53E-32 | √ | √ |
| *Plk1* | -2.1 | 1.91E-17 | X | √ |
| *Set* | -1.2 | 1.30E-36 | √ | √ |
| *Btrc* | -1.2 | 1.19E-18 | √ | √ |
| *Cops6* | -0.76 | 1.23E-11 | √ | √ |
| *Usp37* | -0.52 | 5.07E-06 | √ | √ |
| *Usp22* | -0.47 | 4.31E-06 | √ | √ |

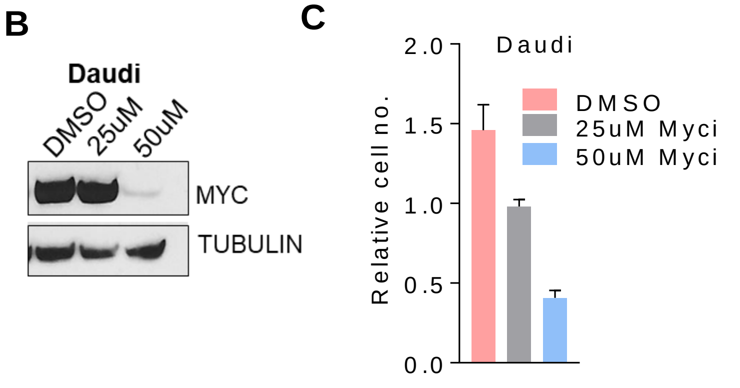


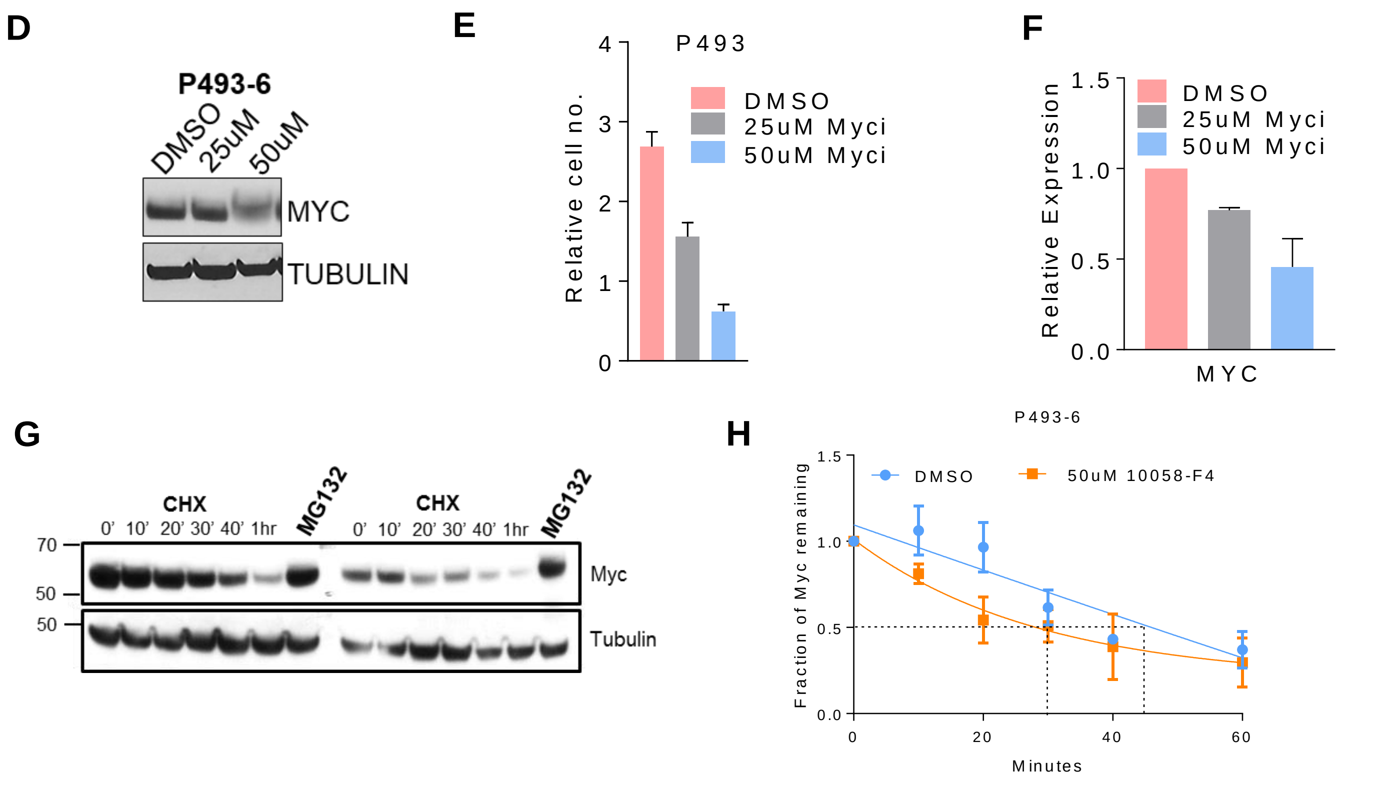


**Supplemental Fig S5 Factors mediating MYC degradation in the absence of MAX**

(A) Expression and ChIP binding data for MYC stability regulating genes in Eμ-*Myc* premalignant cells (ChIP data from Sabo et al, 2014) (B) MYC levels in Daudi cells treated with Myci (C) Relative growth in Myci treated Daudi cells (D) MYC levels in P493-6 cells treated with Myci (E) Cell growth in Myci treated P493-6 cells (F) Relative MYC mRNA levels in Myci treated Daudi cells (G) Representative immunoblot and (H) half-life determination for CHX chase in DMSO and Myci treated P493-6 cells.

### Supplemental Fig. S6

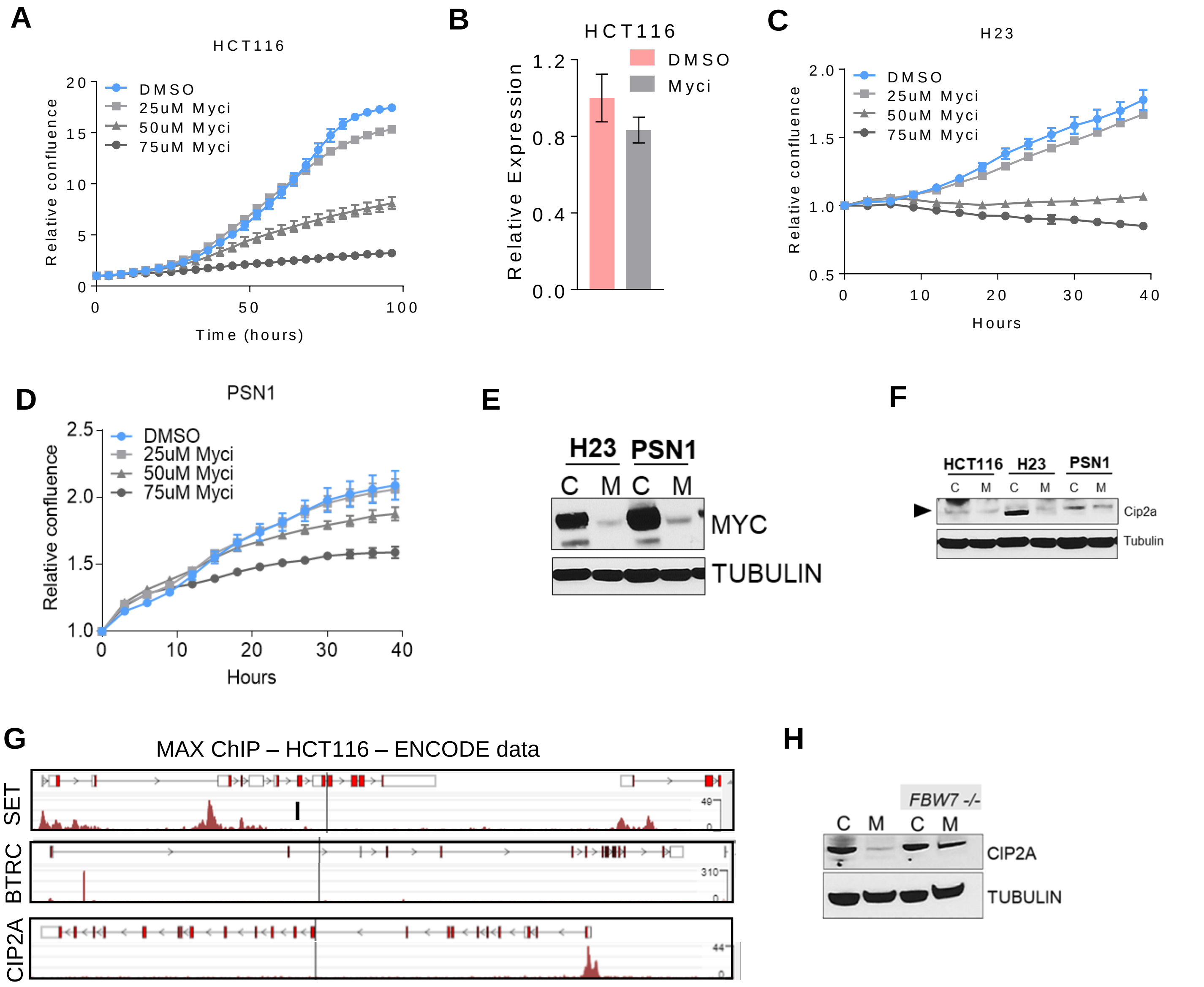


**Supplemental Fig S6 MYC stability is deregulated in a wide spectrum of tumors upon loss of MYC-MAX dimerization.** (A) Cell growth and (B) relative *Myc* mRNA levels in Myci treated HCT116 cells. Growth kinetics of (C) NCI-H23 and (D) PSN-1 cells at different concentrations of Myci. (E) MYC levels in H23 and PSN-1 cells (Control lanes marked C; Myci treated marked M). (F) CIP2A protein levels decrease upon Myci treatment in all 3 cell lines. (G) ENCODE data showing MAX binding at *Cip2a*, *Btrc* and *Set* in HCT116 cells. (H) CIP2A protein levels in HCT FBW7 -/- Myci treated cells

### Supplemental Fig. S7

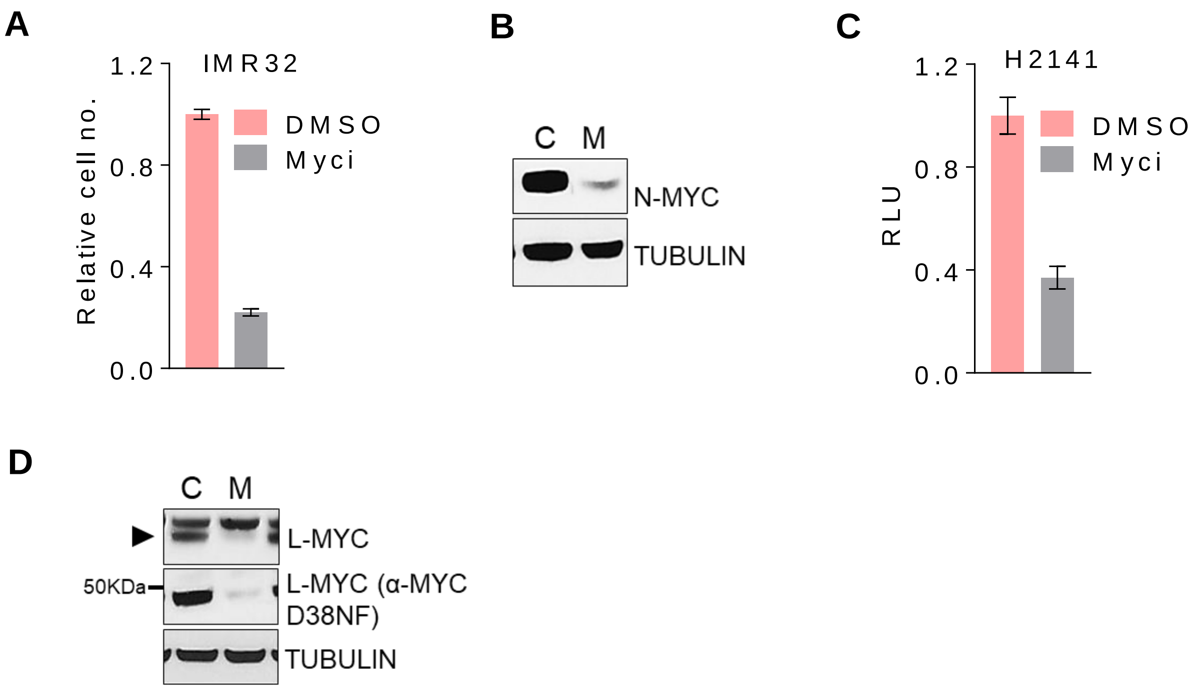


**Supplemental Fig S7 MYC paralog stability is also affected upon disruption of MYC-MAX dimerization** (A) Cell growth and (B) MYCN levels in IMR-32 neuroblastoma cells treated with DMSO or 50uM Myci. (C) Relative cell growth and (D) MYCL levels in Myci vs. control treated L-MYC amplified small cell lung carcinoma line NCI-H2141.

### Supplemental Table s1

| Target | Conjugate | Company | Catalog no. | Purpose |
|---|---|---|---|---|
| B220 | PerCP | BD BIOSCIENCE | 561086 | Flow cytometry |
| CD19 | PE | AFFYMETRIX | 12-0193-82 | Flow cytometry |
| CD21 | FITC | BIOLEGEND | 123407 | Flow cytometry |
| CD23 | APC-Cy7 | BIOLEGEND | 101629 | Flow cytometry |
| IgM | PE-Cy7 | AFFYMETRIX | 25-5790-81 | Flow cytometry |
| IgD | APC-eFLUOR® 780 | AFFYMETRIX | 47-5993-80 | Flow cytometry |
| CD11b | PE | BD BIOSCIENCE | 553311 | Flow cytometry |
| CD3 | FITC | AFFYMETRIX | 11-0031-81 | Flow cytometry |
| CD4 | FITC | AFFYMETRIX | 11-0043-82 | Flow cytometry |
| CD8 | APC-eFLUOR® 780 | AFFYMETRIX | 47-0081-80 | Flow cytometry |
| B220 | purified | BIOLEGEND | 103201 | IF-paraffin |
| Ki67 | purified | ABCAM | ab16667 | IF-paraffin |
| MAX | purified | SANTA CRUZ | sc-197 | IF-paraffin/ Western |
| MYC | purified | CELL SIGNALING | 13987 | Western |
| CIP2A | purified | NOVUS BIOLOGICALS | NB110-59722 | Western |
| total Akt | purified | CELL SIGNALING | 4691 | Western |
| p-Akt Ser 473 | purified | CELL SIGNALING | 4060 | Western |
| p-GSK3b | purified | CELL SIGNALING | 9323S | Western |
| pan GSK3B | purified | SANTA CRUZ | sc-9166 | Western |
| Tubulin | purified | SIGMA | T5326 | Western |
| Actin | purified | SIGMA | A5441 | Western |
| N-MYC | purified | SANTA CRUZ | sc-56729 | Western |
| L-MYC | purified | developed at Fred Hutch | | Western |
| Cleaved caspase 3 | purified | CELL SIGNALING | 9664 | Western |
| RB (total and phospho) | purified | BD BIOSCIENCE | 554136 | Western |

**Supplemental Table s1** Antibodies used in study

### Supplemental Table s2

| Target | Forward | Reverse | Species |
|---|---|---|---|
| Myc | Taqman gene expression assay | Taqman gene expression assay | Mouse |
| Max | Taqman gene expression assay | Taqman gene expression assay | Mouse |
| Rps16 (housekeeping) | Taqman gene expression assay | Taqman gene expression assay | Mouse |
| MYC | CTGAGGAGGAACAAGAAGATGAGGAAG | GTGGGCTGTGAGGAGGTTTGC | Human |
| MAX | TTGTGAATCTGAACTGCTCTACT | CGACTCTGTGCTGCGAAT | Human |
| CIP2A | AGTCAGTACAAAGCCGTGAAG | ATAGTCGTGTGAGTTTCTGTCC | Human |
| BTRC | CTTAAATGGACACAAACGAGGC | CAACGCACCAATTCCTCATG | Human |
| SET | AAATATAACAAACTCCGCCAACC | CAGTGCCTCTTCATCTTCCTC | Human |
| PLK1 | ACAGTTTCGAGGTGGATGTG | GGTTGATGTGCTTGGGAATAC | Human |
| GUSB (housekeeping) | CCTGCGTGTCCCTTCCTC | CGTTCTGGTCTGCCGTGAA | Human |

**Supplemental Table s2** Real time primers used in study
